# Human *in vitro* model of material-driven vascular regeneration reveals how cyclic stretch and shear stress differentially modulate inflammation and tissue formation

**DOI:** 10.1101/755157

**Authors:** Eline E. van Haaften, Tamar B. Wissing, Nicholas A. Kurniawan, Anthal I.P.M. Smits, Carlijn V.C. Bouten

**Author notes:** E.E. van Haaften MSc, Dr. T.B. Wissing, Dr. N.A. Kurniawan, Dr. A.I.P.M. Smits, Prof. dr. C.V.C. Bouten, P.O. Box 513, 5600 MB Eindhoven, the Netherlands.

## Abstract

Resorbable synthetic scaffolds designed to regenerate living tissues and organs inside the body emerge as a clinically attractive technology to replace diseased blood vessels. However, mismatches between scaffold design and *in vivo* hemodynamic loading (i.e., cyclic stretch and shear stress) can result in aberrant inflammation and adverse tissue remodeling, leading to premature graft failure. Yet, the underlying mechanisms remain elusive. Here, a human *in vitro* model is presented that mimics the transient local inflammatory and biomechanical environments that drive scaffold-guided tissue regeneration. The model is based on the co-culture of human (myo)fibroblasts and macrophages in a bioreactor platform that decouples cyclic stretch and shear stress. Using a resorbable supramolecular elastomer as the scaffold material, it is revealed that cyclic stretch initially reduces pro-inflammatory cytokine secretion and, especially when combined with shear stress, stimulates IL-10 secretion. Moreover, cyclic stretch stimulates downstream (myo)fibroblast proliferation and neotissue formation. In turn, shear stress attenuates cyclic-stretch-induced tissue growth by enhancing MMP-1/TIMP-1-mediated collagen remodeling, and synergistically alters (myo)fibroblast phenotype when combined with cyclic stretch. The findings suggest that shear stress acts as a stabilizing factor in cyclic stretch-induced tissue formation and highlight the distinct roles of hemodynamic loads in the design of resorbable vascular grafts.

## 2 Introduction

The use of bioresorbable replacements (e.g., cell-free scaffolds) designed to direct tissue regeneration at the locus of implantation has emerged as a promising strategy to create replacements that are living and adaptive.^[1][2]^ This technology has the potential to not only reduce the clinical demand for autologous living blood vessel replacements but also bypass the challenges associated with synthetic, nondegradable vascular substitutes. The regenerative response is initiated by the infiltration of immune cells, e.g., macrophages, followed by the attraction, migration, and distribution of tissue-producing cells that deposit *de novo* extracellular matrix (ECM) in the first weeks after implantation (**Figure 1A**).^[1]^ Ideally, the deposited tissue is remodeled, guided by physiological hemodynamic loads, to resemble a tissue with native-like structural and functional properties, while the scaffold slowly degrades.^[3][4]^

**Figure 1.**
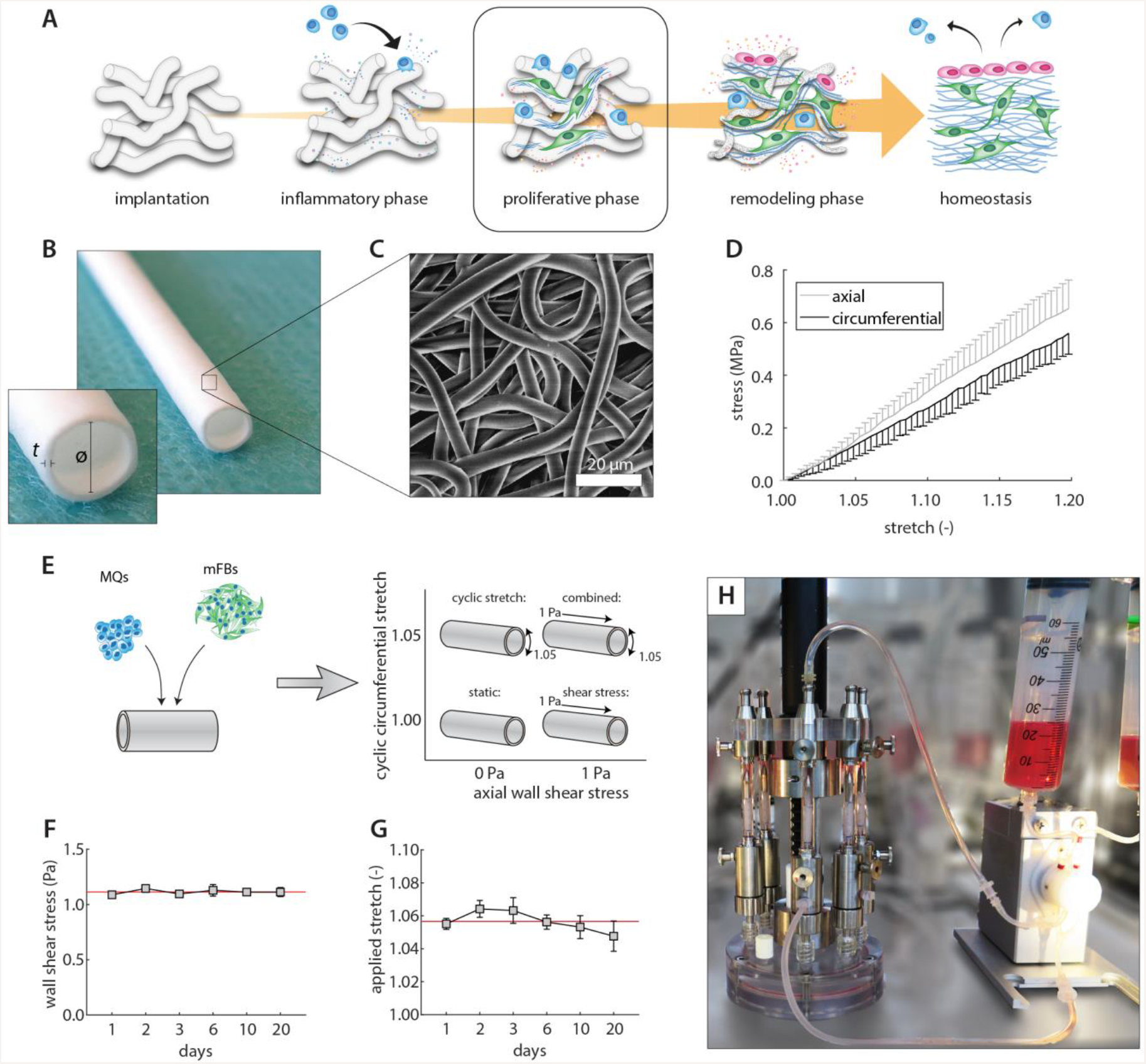
*In vitro* set-up of the inflammatory response and biomechanical environment during scaffold-guided tissue regeneration. A) The different phases of scaffold-driven tissue regeneration at the host’s functional site. Our model focusses on the proliferative phase, in which macrophages and tissue-producing cells have colonized the scaffold material. B) Electrospun PCL-BU vascular scaffold is selected as a biomaterial (diameter (ø) and thickness (t) are indicated). C) Scanning electron microscopy image of the scaffold microstructure. D) Averaged stress-stretch curve (*n* = 4) of the bare scaffold material prior to the experiment. E) Scaffolds containing macrophages (MQs) and (myo)fibroblasts (mFBs) are exposed to different loading conditions and compared with static controls. F-G) The monitored wall shear stresses and cyclic circumferential stretches during the experiment. H) Cyclic stretch, shear stress, or a combination of both loads is imposed on the scaffold constructs using a bioreactor platform. Panels A and H are reproduced with permission.^[1][33]^

Despite encouraging *in vivo* proof-of-concept studies,^[5][6]^ early stenosis and aneurysm formation remain prevalent complications in *in situ* vascular tissue engineering.^[7]–[9]^ These complications are often attributed to a mismatch between scaffold properties (e.g., microstructure, geometry, and mechanical properties) and mechanical loading, resulting in aberrant inflammation and pathological tissue deposition. It is therefore important to attain a more fundamental understanding of the cell and tissue responses to the chemo-physical micro-environment during *in situ* tissue regeneration.

One major process that plays a critical role for successful *in situ* regeneration is the host immune response. While the immune response is essential for the initiation and guidance of wound healing, it can also contribute to adverse tissue deposition (e.g., fibrosis) and scaffold failure (e.g., accelerated degradation) in conditions of chronic inflammation.^[1]^ Particularly, due to its phenotypic and functional plasticity, the monocyte-derived macrophage has been identified as the commanding cellular player in the initial immune response, driving biomaterial degradation as well as tissue formation and remodeling.^[1][10]–[16]^ The macrophage “prepares the ground”—by secreting cytokines and degrading the scaffold—for colonizing cells (e.g., infiltrating mature fibroblasts, smooth muscle cells, as well as stem/progenitor cells) to produce a new tissue.^[17]^ Thus, the interplay between macrophages and tissue-producing cells within the scaffold environment will largely determine the functionality of the resulting neotissue.

Co-cultures of macrophages and tissue-producing cells revealed that paracrine signaling between these cell types promote cellular recruitment, attachment, migration, and distribution throughout porous scaffolds.^[10]–[12][14]^ Moreover, paracrine factors secreted by different macrophage phenotypes differentially regulate fibroblast and smooth muscle cell behavior (e.g., proliferation and phenotype).^[13][15]^ Multiple macrophage phenotypes have been identified, spanning a spectrum with the classical pro-inflammatory (M1) and the alternative anti-inflammatory (M2) phenotypes at the extremes.^[18]^ Paracrine factors secreted by M2 macrophages (e.g., TGF-β, PDGF) are known to stimulate fibroblast proliferation and collagen formation, whereas factors secreted by M1 macrophages (e.g., TNF-α, MMP-7) give rise to a more pro-inflammatory fibroblast with tissue degrading as well as pro-fibrotic properties.^[13][15]^ Of note, fibroblast behavior can be reversed by reversing or removing the paracrine signals, emphasizing fibroblast plasticity.^[13]^

The phenotypic state of the macrophage is ambiguous, often displaying both pro- and anti-inflammatory features simultaneously, and depends on the phase of tissue repair and the biochemical cues from the cellular environment it is exposed to.^[19]–[21]^ In addition, while the exact (intra)cellular mechanisms remain elusive, it is increasingly acknowledged that macrophages are mechanosensitive and adjust their function in response to both physical and mechanical cues.^[22]–[24]^ For example, it has been reported that lower strains (8%) permit macrophage polarization towards a reparative M2 profile in 3-dimensional (3D) electrospun scaffolds, whereas higher strains (12%) promote a more pro-inflammatory M1 profile, both in terms of cell phenotype and cytokine secretion.^[25][26]^ Moreover, Battiston et al. demonstrated increased collagen deposition in cyclically-stretched scaffolds seeded with monocytes and SMCs.^[10]^ In addition to mechanical stretching, the relation between flow-induced shear stress and cell behavior has been the topic of multiple studies.^[27]–[30]^ These studies have provided insights into the effect of either shear stress or cyclic stretch on various cell types individually, especially on 2D substrates.^[31][32]^ However, knowledge on the relative contribution of shear stress and cyclic stretch to tissue regeneration in a more physiologically relevant 3D environment remains unknown.

In the present study, motivated to improve the long-term clinical performance of *in situ* tissue engineered vascular grafts, we employ a human *in vitro* model to identify the mutual roles of physiological levels of shear stress and cyclic stretch on macrophage/fibroblast-mediated neotissue formation. Using our recently developed bioreactor,^[33]^ we reveal that these loads differently regulate immune profiles, tissue growth, and matrix remodeling in co-cultures of (myo)fibroblasts and monocytes in electrospun vascular polycaprolactone bis-urea (PCL-BU) scaffolds. Whereas shear stress abrogates stretch-induced tissue growth and stimulates collageneous remodeling, cyclic stretch inhibits shear stress-driven secretion of pro-inflammatory cytokines (MCP-1 and IL-6). Moreover, both loads cumulatively affect anti-inflammatory IL-10 secretion and (myo)fibroblast phenotype. Together, the data highlight the importance of considering the effects of both cyclic stretch and shear stress during the different phases of *in situ* vascular regeneration.

## 3 Results & Discussion

### 3.1 Characterization of scaffolds and hemodynamic loads

PCL-BU was chosen as a biomaterial as this supramolecular elastomer has shown its potential for *in situ* cardiovascular TE applications.^[34][35]^ PCL-BU was processed using electrospinning into vascular scaffolds with an inner diameter of 3 mm and wall thickness of 200 μm (Figure 1B). The resulting scaffolds, with a fiber diameter of 4.58 ± 0.34 μm, showed a highly porous and isotropic microstructure, allowing for cell infiltration (Figure 1C). Biaxial tensile testing revealed that the material behaves linearly (Figure 1D) within the applied loading regime (up to 1.08 stretch), but is slightly stiffer in the axial direction (3.6 ± 0.9 MPa) in comparison to in the circumferential direction (3.0 ± 0.4 MPa). The scaffolds were seeded with a 2:1 mixture of human primary monocytes and (myo)fibroblasts^[10][11]^ using fibrin as a cell carrier (Figure 1E), and cultured under healthy physiological levels of shear stress^[36]^ (1.11 ± 0.08 Pa, Figure 1F), cyclic stretch^[2]^ (1.06 ± 0.02, Figure 1G), or a combination of both for 20 days in our bioreactor (Figure 1H). Since the scaffold constructs were mounted around impermeable silicone tubing for the application of circumferential stretch, transmural flows are assumed to be negligible.

### 3.2 Cyclic stretch promotes (myo)fibroblast proliferation

We first assessed whether co-culture of macrophages and (myo)fibroblasts for 20 days in the vascular scaffolds resulted in tissue deposition. Indeed, SEM analysis revealed an increasing presence of cells and ECM deposition in the scaffolds from day 3 to day 20 in all sample groups (**Figure 2A**). Elevated DNA levels were found in the presence of cyclic stretch at day 20 (Figure 2B), confirming that cyclic stretch stimulates cell proliferation.^[37]–[40]^ In contrast, shear stress alone was not found to lead to increased DNA levels. To identify which cell population is responsible for this overall proliferation, we stained the cells with KI67. We found that KI67 co-localized only with vimentin-positive (myo)fibroblasts, which adopted an elongated spindle-shaped morphology (Figure 2C). The macrophages (CD45-positive cells in Figure 2C) did not proliferate but remained viable with a more rounded morphology throughout the duration of the experiment. Overall, these results show a stimulatory effect of cyclic stretch on early (myo)fibroblast proliferation, irrespective of the presence of shear stress.

**Figure 2.**
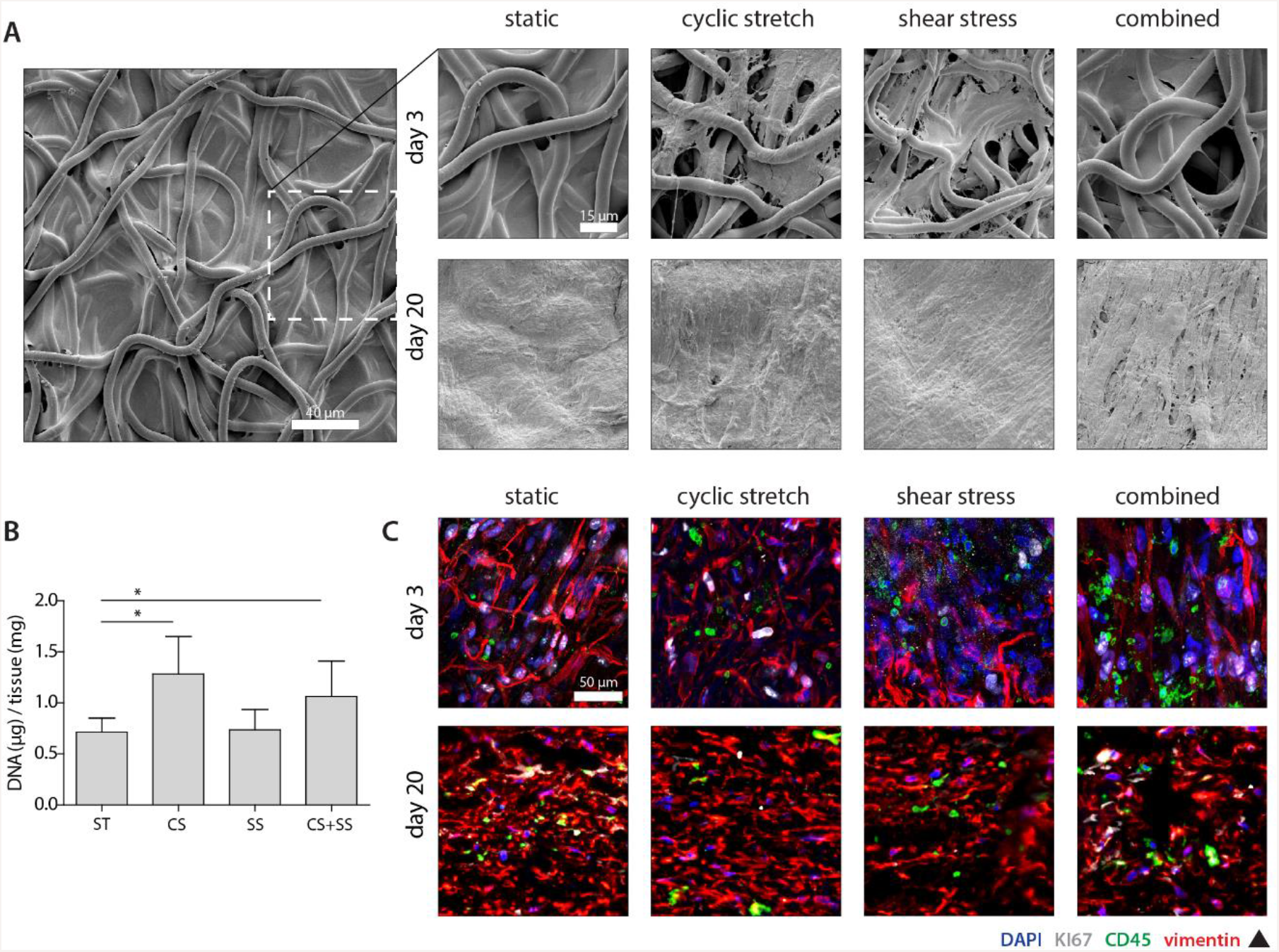
Cell and tissue growth under cyclic stretch and shear stress. A) Overall cell and tissue morphology at day 3 (*top*; low magnification of the static condition is shown *left*) and day 20 (*bottom*). B) DNA content at day 20 (*n* ≥ 5 per group) normalized to the total construct mass (* *p* < 0.05). C) (Myo)fibroblast proliferation at day 3 (*top*, whole-mounts) and day 20 (*bottom*, cross-sections) (blue = DAPI; white = KI67; green = CD45; red = vimentin). Abbreviations: static (ST), cyclic stretch (CS), SS (shear stress).

### 3.3 Loading regimes differentially affect cell inflammatory response and phenotype

To characterize the inflammatory environment in the vascular constructs under different regimes of hemodynamic loading, we examined the cytokine secretion profiles in the medium (**Table S1**). No obvious overall difference between different loading regimes at day 3 was observed, except for the secretion of MMP-9, which was clearly enhanced with cyclic stretch (**Figure 3A**). All cytokine secretion levels increased with longer culture, except for the pro-inflammatory protein TNF-α and the anti-inflammatory IL-10. More specifically, TNF-α was present in all mechanically loaded samples at day 3 but not at day 20, and IL-10 levels slightly decreased with time, but remained stable and was highest when both loads were combined. However, it should be noted that the concentrations of these two cytokines were relatively low compared to other cytokines.

**Figure 3.**
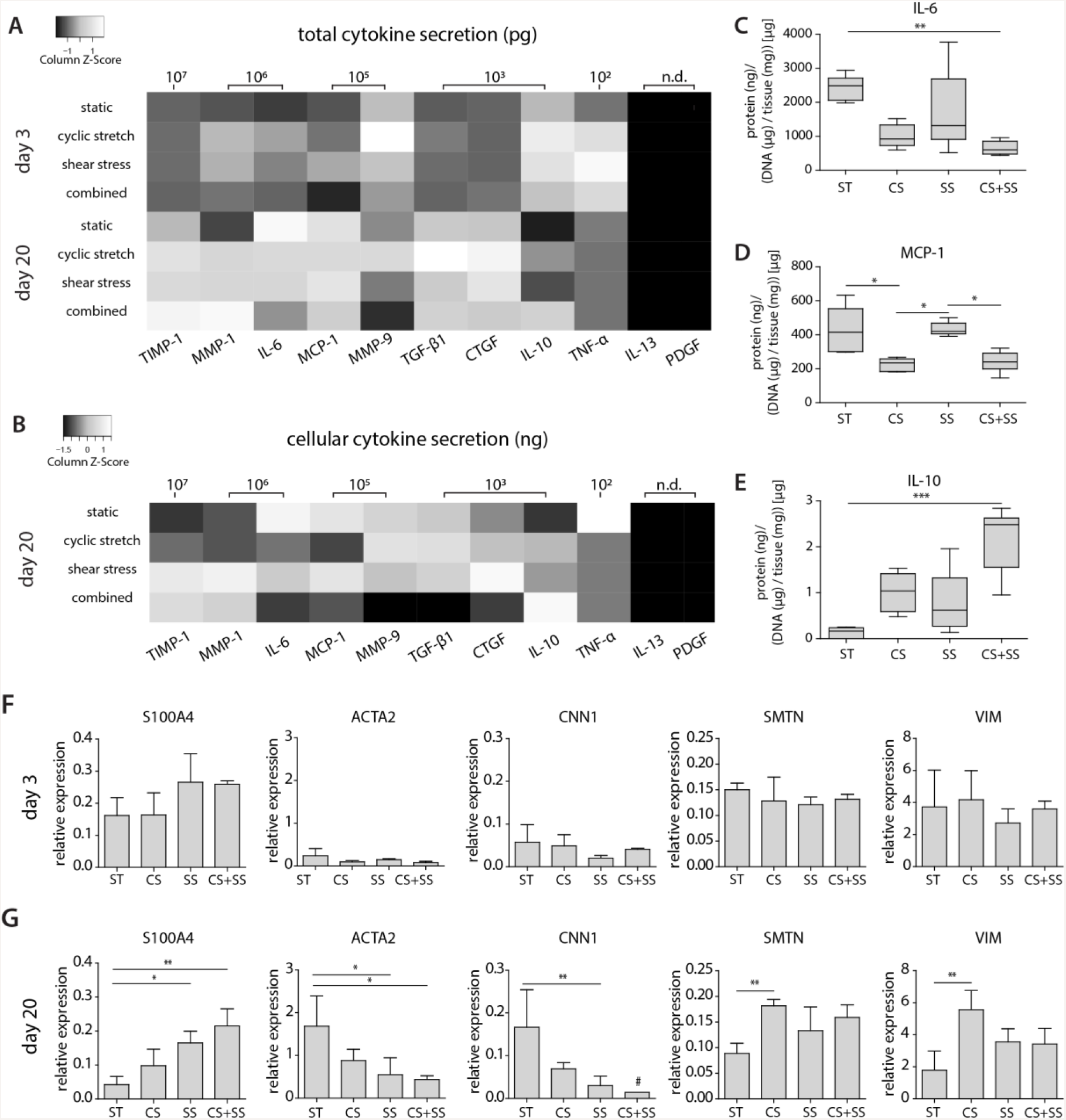
Inflammatory environment and (myo)fibroblast phenotype in the vascular construct. Heat maps of the A) total cytokine secretion at day 3 (*n* = 4 per group) and day 20 (*n* ≥ 5 per group) and B) cytokine secretion at day 20 normalized to DNA content (gray scales are normalized for each cytokine; TNF-α was only detected in the mechanical loaded groups at day 3 (*n* ≥ 2); n.d., not detected). C-E) Boxplots for a selection of cytokines at day 20 normalized to DNA content (see **Figure S1**A, B and **Figure S2** for the boxplots of all cytokines). F-G) Relative gene expressions of (myo)fibroblast-specific phenotypic markers at day 3 (*n* ≥ 3 per group, except for calponin (*n* = 2 in combined group)) and day 20 (*n* ≥ 4 per group, except for calponin *n* = 3 in cyclic stretch group)). # measured at or below the detection limit. (* *p* < 0.05, ** *p* < 0.01, *** *p* < 0.001). Abbreviations: metallopeptidase inhibitor (TIMP), matrix metalloproteinase (MMP), interleukin (IL), monocyte chemoattractant protein 1 (MCP-1), transforming growth factor beta 1 (TGF-β1), connective tissue growth factor (CTGF), tumor necrosis factor alpha (TNF-α), platelet derived growth factor (PDGF). S100 calcium binding protein A4 (S100A4), Alpha Smooth Muscle Actin (ACTA2), calponin 1 (CNN1), smoothelin (SMTN), vimentin (VIM), static (ST), cyclic stretch (CS), SS (shear stress).

To examine in more detail the cellular response to the applied hemodynamic loading, we normalized the total protein secretion levels to the DNA content at day 20 (Figure 3B). Cyclic stretch contributed to a clear reduction of pro-inflammatory MCP-1 and IL-6, together with a concomitant upregulation of the anti-inflammatory protein IL-10, especially when combined with shear stress (Figures 3C-E).^[41]^ These changes strongly suggest that cyclic stretching tunes and prevents excessive inflammation by promoting an anti-inflammatory environment. This finding corroborates with previous studies that showed that moderate stretches can promote an anti-inflammatory environment, both via direct mechanosensing and via indirect paracrine signaling.^[25][42][43]^ Furthermore, we found that the secretion of TGF-β1 and CTGF was reduced with combined loads, implying a suppressed tissue-stimulatory environment (e.g., to produce collagen) compared to no or single loads. In contrast, MMP-1 and TIMP-1 secretion was increased with combined loads, implying an enhanced tissue-remodeling environment (e.g., to remodel collagen).

We next examined whether these loading-dependent inflammatory responses modulate the (myo)fibroblast phenotype. Differences in the gene expressions representing (myo)fibroblast phenotypic markers became most apparent at day 20 (Figure 3F, G). Especially for αSMA, a strong increase in expression was observed from day 3 to day 20. Interestingly, at day 20, both cyclic stretch and shear stress application contributed to an upregulation of S100A4, a marker for activated fibroblasts that has been recently related to inflammation,^[44][45]^ and a downregulation of the contractile markers αSMA and calponin in a synergistic manner, suggesting that mechanical loading, in either or combined form, reduces cell contractility (Figure 3G). Nevertheless, the expression of smoothelin, another specific marker related to cell contractility, as well as vimentin, were significantly enhanced in the presence of cyclic stretch and to a lower extent in the presence of shear stress.

Both calponin and αSMA have been found to be regulated via cytokines (e.g., MCP-1 and IL-6^[12]^) as well as loading regime (e.g., shear stress^[46]–[48]^). As such, we detect paradoxical load-dependent changes in phenotype marker expressions that are likely to be a combined effect of direct mechanotransduction (i.e., (myo)fibroblast response to cyclic stretch and superficial/interstitial shear stress) and the indirect, cell-mediated cytokine profiles in the medium.

### 3.4 Shear stress dampens cyclic-stretch-induced tissue formation and enhances MMP-1/TIMP-1-mediated remodeling

Cyclic stretch, depending on the magnitude and duration, is known to stimulate matrix formation.^[37][38][49][50]^ Since the (myo)fibroblast phenotype was affected by the applied loading regime, we hypothesized that the loads applied in this study also affect vascular regeneration by modulating tissue growth and remodeling. We found that, in general, the gene expression of growth and remodeling markers increased over time, as indicated by the brighter colors in the heat maps at day 20 compared to day 3 (**Figure 4A**). Most strikingly, at day 20, cyclic stretch stimulated the expression of several markers related to the production of collageneous matrix (collagen I, collagen III), elastic matrix (elastin, fibrillin 1), and GAGs (versican), but not in the presence of shear stress (Figure 4A, B). Thus, shear stress seemed to have a dampening effect on stretch-induced matrix growth. This is particularly intriguing because shear stress has been previously suggested to be a regulating factor in matrix deposition by inhibiting cell proliferation and neointimal thickening under elevated levels of shear stress.^[51]^

**Figure 4.**
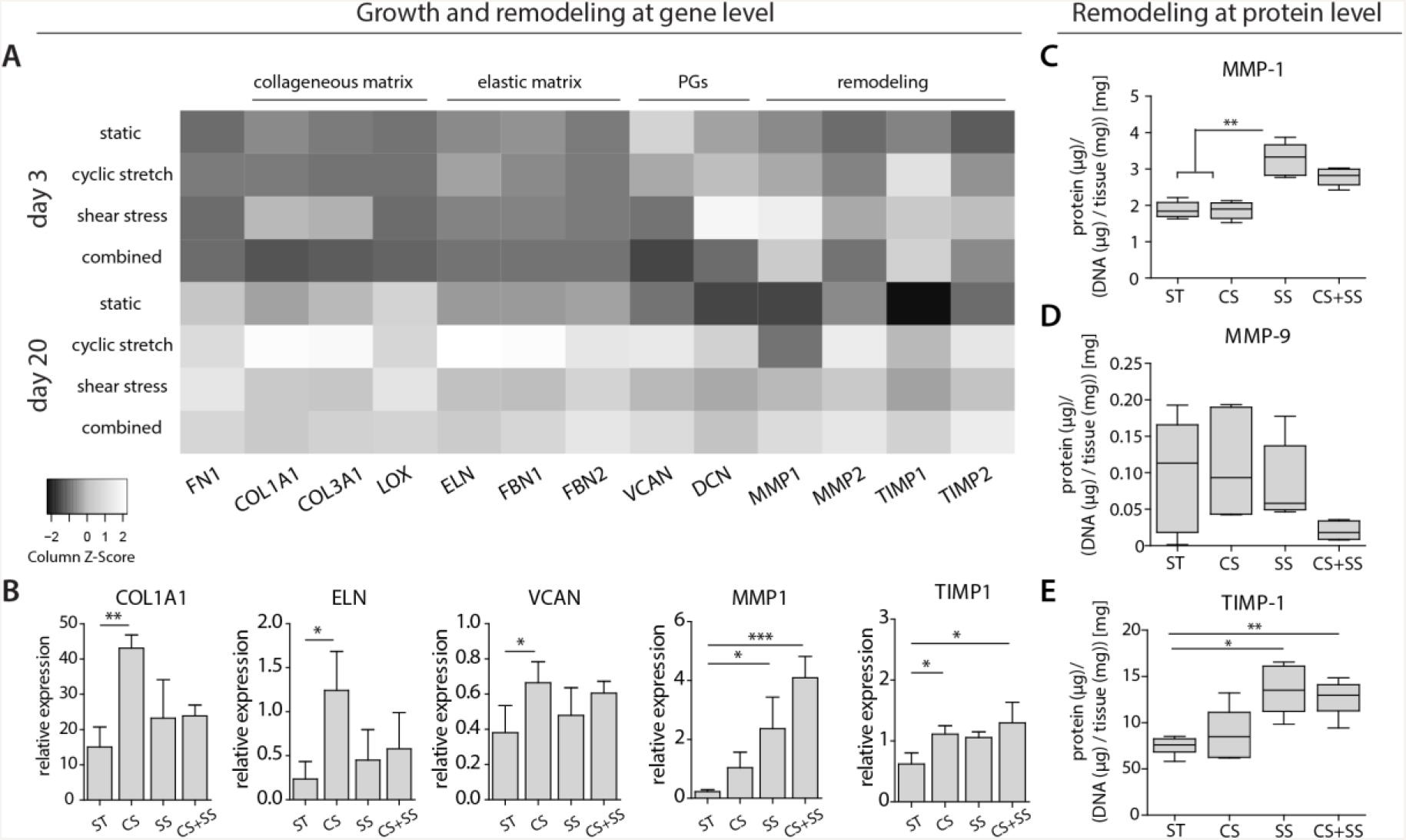
Changes to the markers of tissue growth and remodeling in response to hemodynamic loading. A) Heat maps of relative expression (normalized to GAPDH) for genes related to collageneous matrix, elastic matrix, proteoglycans (PGs), and remodeling at day 3 and day 20 (gray scales are normalized for each gene). *n* ≥ 3 per group (day 3) and *n* ≥ 4 per group (day 20), except for LOX at day 3 (combined, *n* = 2) and FBN2 at day 20 (static, *n* = 3 and cyclic stretch, *n* = 2). B) Expression of a selection of genes at day 20 (see **Figure S3** for the expression levels of all genes at day 3 and day 20). C-E) Protein expressions of matrix metalloproteinases (MMPs) and tissue inhibitors of MMPs (TIMPs) (normalized to DNA content, *n* ≥ 5 per group) at day 20 (see Figure S1A, B for the boxplots of the uncorrected protein expression at day 3 and day 20). * *p* < 0.05, ** *p* < 0.01, *** *p* < 0.001. Abbreviations: static (ST), CS (cyclic stretch), SS (shear stress), fibronectin (FN1), collagen I (COL1A1), collagen III (COL3A1), lysyl oxidase (LOX), elastin (ELN), fibrillin (FBN), verscian (VCAN), decorin (DCN), matrix metalloproteinase (MMP), metallopeptidase inhibitor (TIMP), static (ST), cyclic stretch (CS), SS (shear stress).

In terms of tissue remodeling, we found elevated levels of gene and protein expressions of MMPs and TIMPs upon loading compared to the static controls at day 20. Whereas shear stress stimulated MMP-1 and TIMP-1 gene expression and protein secretion, cyclic stretch independently stimulated the expression of MMP-2 and TIMP-2. MMP-9, on the other hand, was only minimally secreted and no clear differences between loading conditions were found (Figure 4A, C-E). Together, these observations suggest that there is more effective tissue remodeling when both loads are applied. Although comparable observations in literature are found on gene and protein levels when either shear stress or stretch is applied,^[52]–[57]^ these data should be interpreted with caution as they represent only snapshots of the remodeling process without discriminating between latent and active forms of the proteases. Further insight into the activity of the secreted proteases can be provided by zymography,^[58]^ which we additionally performed to test for MMP-2 and MMP-9 activity (i.e., using gelatin zymography, see Figure S1C-E). These data confirm that the secreted proteases are active and thus able to remodel the newly formed matrix.

### 3.5 Shear stress abrogates cyclic-stretch-induced matrix stiffening

The long-term clinical performance of our resorbable vascular scaffolds is largely determined by the structural and compositional properties of this newly formed matrix. Thus, we next sought to identify the effect of hemodynamic loading on the organization of the neotissue. At the superficial tissue layer (i.e., up to a depth of 25 μm into the newly formed tissue, free from the influence of the scaffold fibers), cyclic stretch and shear stress were found to promote the (myo)fibroblasts and collagen to align in a right-handed helical structure with an angle of around 60° with respect to the circumferential axis (**Figure 5A-D, G**). To make a direct comparison across all loading conditions, we plotted the average collagen and actin distributions in a single figure (Figure 5E, F). Cyclic stretch was found to result in a strong alignment of the actin fibers, and to a lesser extent in the presence of shear stress (Figure 5F). A similar but less pronounced effect was found in the collagen structures (Figure 5E). In contrast, inside the scaffold, cells and matrix organization consistently followed the scaffold fibers (**Figure S4**). In line with the earlier findings of de Jonge et al.,^[3]^ these results demonstrate the importance of the contact-guiding cue provided by the micro-fibrous scaffold and the impact of both loads on late-stage cell and matrix reorientation (i.e., when the scaffold has degraded). The exact reorientation response to the applied loading regime, however, also depends on the tissue constraints, dimensionality (2D vs. 3D), and the magnitude and type (laminar vs. oscillatory) of the loads.^[30][59]–[62]^

**Figure 5.**
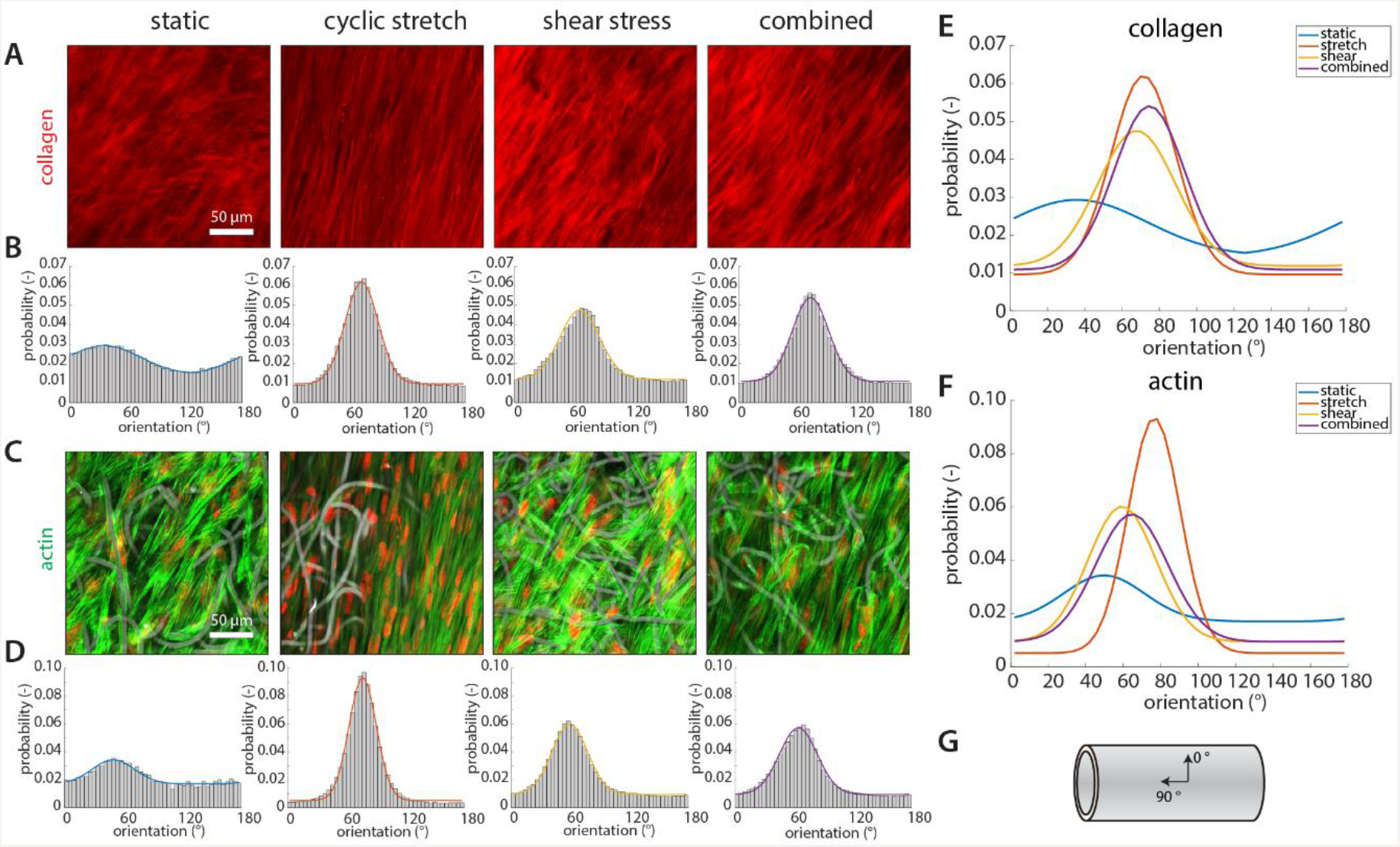
Matrix and cell organization at day 20. A, C) Maximum intensity projection of the collagen organization (A, red = collagen) and the actin fibers (C, green = actin; red = nuclei) at the outer side of the constructs. B, D) Quantification of the collagen and actin fiber distribution from the images (*n* ≥ 5 per group, each solid line is a Gaussian fit with baseline through the data points). E, F) Overlay plot of the quantified collagen and actin fiber orientation distribution for each condition. G) Schematic showing the imaging direction.

It is tempting to speculate that the hemodynamic-loading-dependent organization of the neotissue is responsible for changes in the construct mechanical properties. It should be noted, however, that the superficial tissue layer is likely too thin to markedly affect the mechanical performance of the overall construct. Therefore, we also analyzed the compositional properties of the constructs. Quantification of sample thickness from the tissue sections indicated that the presence of cyclic stretch slightly, but not significantly, led to the formation of a thicker tissue (**Figure 6A**). Collagen staining and HYP quantification further revealed that loading regime influenced collagen composition, both in terms of the type and the amount of the collagen present in the construct (Figure 6B, D-F). Cyclic stretch stimulated the formation of more numerous and thicker fibers of collagen type I, but this effect was overruled by the dampening effect of shear stress, in direct correspondence with our earlier finding at the gene expression level (Figure 4B). In contrast to collagen, the formation of GAGs was independent of the applied loading regime (Figure 6C, G).

**Figure 6.**
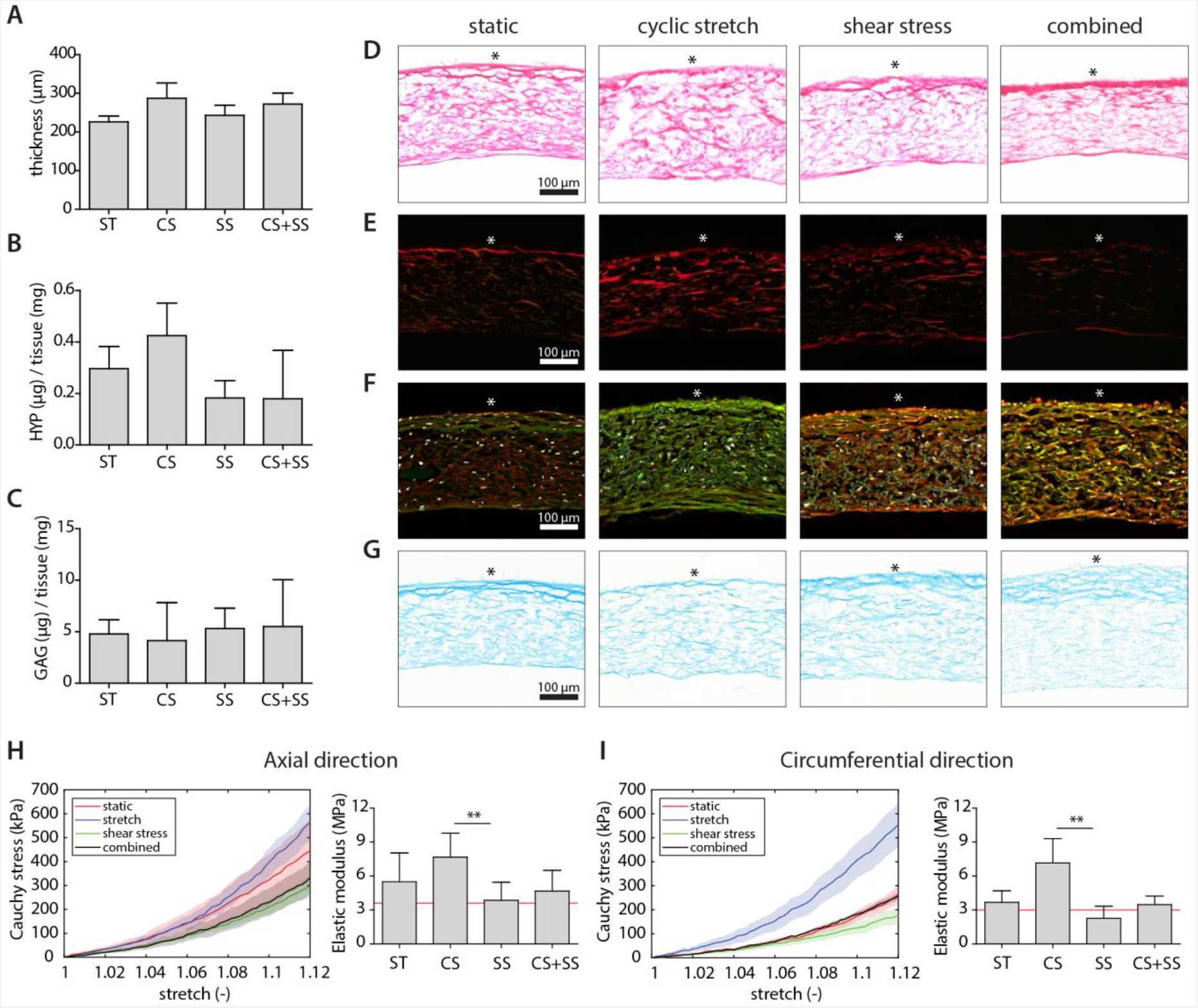
Matrix composition and mechanical properties at day 20. A) Construct thickness (*n* = 5 per group). B) Hydroxyproline and C) glycosaminoglycan levels normalized total construct mass (*n* ≥ 5 per group, see **Figure S5** for the normalization to DNA content). D-G) Cross-sections stained for Picrosirius red under bright field (D) and polarized light (E); collagen type I (green), collagen type III (red), and DAPI (white) (F); and Alcian blue (G) (* denotes the outer side of the constructs). H, I) Averaged stress–stretch curves of the cultured constructs in the axial (H, left) and circumferential (I, left) direction at day 20 (*n* ≥ 3 per group, shaded areas represent the standard error of mean), and quantification of the elastic modulus at 1.10 stretch from the stress–stretch curves in the axial direction (H, right) and the circumferential direction (G, right) (** p < 0.01, the *red line* indicates bare scaffold properties). Abbreviations: static (ST), CS (cyclic stretch), SS (shear stress), hydroxyproline (HYP), glycosaminoglycans (GAGs).

The effects of hemodynamic loading on the tissue growth and remodeling are reflected in the mechanical properties of the constructs, which are important for maintaining vascular integrity during *in situ* regeneration. Biaxial tensile tests revealed that the engineered tissue constructs exhibited some degree of non-linear behavior in both the axial and circumferential directions and were slightly stiffer in the axial direction (Figures 6H, I). This is in contrast to the largely isotropic structure and mechanical properties of the bare scaffolds (3.6 ± 0.9 MPa and 3.0 ± 0.4 MPa in axial and circumferential direction respectively). Cyclic stretch resulted in the stiffest tissue (7.7 ± 2.1 MPa and 7.2 ± 2.2 MPa in Figures 6H and I). Interestingly, the presence of shear stress abrogates this stretch-induced matrix stiffening in both directions, leading to tissue stiffness that is almost indistinguishable from that of the bare scaffold (4.7 ± 1.8 MPa and 3.5 ± 0.8 MPa in Figures 6H and I).

Taken together, we observed more collagen and more predominant collagen type I in the cyclically stretched samples, but negligible differences in lysyl oxidase (LOX) expression (a catalyzer of covalent fiber crosslinking important for the stabilization of collagen and elastin networks) in comparison to the other experimental groups (Figure 4A). Based on these findings, we conclude that the increased axial and circumferential stiffness in the cyclically stretched samples should be attributed to differences in the tissue composition (i.e., quantity and type). Where cyclic stretch stimulated the formation of a stiffer matrix, containing more and predominantly collagen type I, shear stress stimulated the deposition of collagen type III and the secretion of MMP-1 and TIMP-1. Our observations therefore indicate that shear stress plays a key regulating role in the progression of tissue growth and remodeling, especially in the presence of cyclic stretch.

### 3.6 Future perspectives

This study aids in understanding the mechanobiological processes that underlie vascular regeneration. We opted for a more physiological co-culture with the inherent difficulty that the cell ratio varied with time, due to (myo)fibroblast proliferation and a reduction in the macrophage number (e.g., as a result of apoptosis and potential transdifferentiation). From a fundamental point-of-view, it would be interesting to discriminate between the contributions of each individual cell type (i.e., myofibroblasts and macrophage) and explore potential donor variability using multiple donors. For the study of separate cell populations, the reader is referred to Wissing et al. (2019b, *under review*).

The (myo)fibroblasts that were used in our model are a consistent vascular cell source known to play a considerable role in tissue deposition and remodeling, with the ability to accommodate to extreme loading conditions.^[63][64]^ Nevertheless, as these cells originate from the venous circulation, they may have been ‘overstressed’ as they are not ‘primed’ to arterial loading conditions.^[36][65]^ Potential overstressing of the cells could have contributed to the more synthetic phenotype that we observed in the presence of hemodynamic loads, with only subtle differences between the applied loading regimes. It will therefore be instructive to repeat the experiments with human cells from various origins (e.g., arteries, bone marrow).

## 4 Conclusion

In summary, by employing a dynamic 3D human *in vitro* model, we successfully discriminated for the first time the distinct and combined effects of cyclic stretch and shear stress during material-driven vascular tissue regeneration (**Scheme 1**). Most importantly, during the early inflammatory and proliferative phase, shear stress can be recognized as a stabilizing factor in cyclic stretch-induced tissue formation, highlighting that both hemodynamic loads should be considered in the design of resorbable vascular grafts. Our approach paves the way for future research, in which this study should be extended to unravel the underlying principles of late-stage vascular tissue homeostasis, for example by focusing on the individual and combined responses of cells (e.g., immune and progenitor cells) to variable hemodynamic loading regimes, as well as the changes that occur after endothelialization. This knowledge is critical to identify optimal scaffold designs that improve the long-term clinical performance of *in situ* tissue engineered blood vessels.

**Scheme 1.**
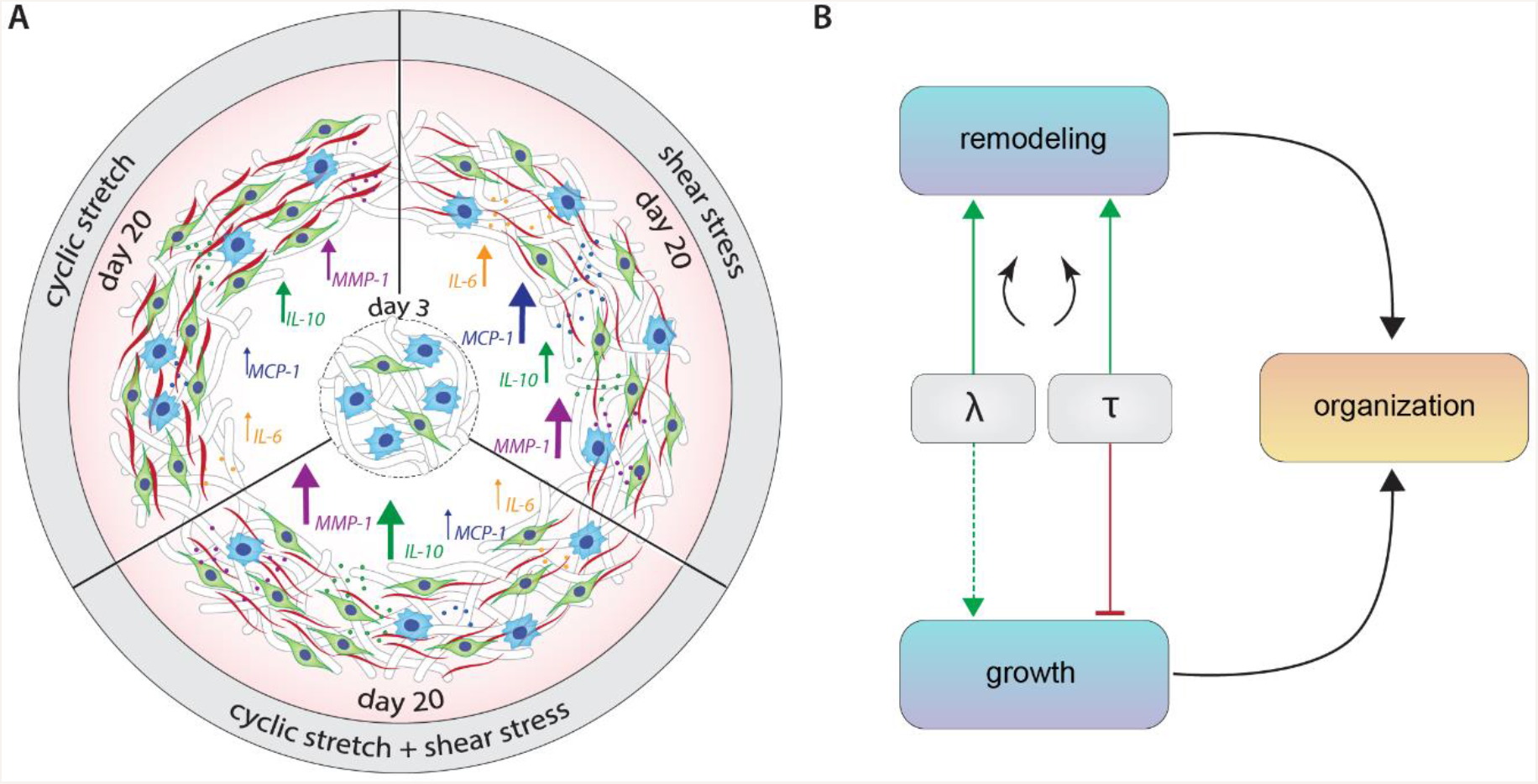
Graphical synopsis of the main findings of the study. A) Cyclic stretch stimulates matrix growth and cell proliferation, and inhibits shear stress-driven secretion of pro-inflammatory cytokines (MCP-1 and IL-6), while both loads contribute to the secretion of MMP-1 and anti-inflammatory IL-10. B) At the tissue level, shear stress (τ) abrogates stretch-induced tissue growth and stimulates, especially in the presence of cyclic stretch (λ), collageneous remodeling, resulting in a tissue with directed collagen fibers.

## 5 Experimental Section

### Scaffold preparation

Isotropic 3 mm (inner diameter) scaffolds were electrospun from polymer solutions containing 15 % (w/w) bis-urea-modified poly-ε-caprolactone (PCL-BU, SyMO-Chem, Eindhoven, The Netherlands) and 85 % (w/w) chloroform (Sigma; 372978). The solutions were delivered at 40 μL min^−1^ via a charged needle (16 kV) on a negatively charged, rotating mandrel (−1 kV, 500 rpm, 3 mm diameter) in a climate-controlled cabinet (23 °C and 30 % humidity, IME Technologies, Geldrop, The Netherlands). The distance between the needle and the mandrel was kept constant at 16 cm. The resulting scaffolds were dried overnight under vacuum to remove solvent remnants and placed over silicone tubing (2.8 mm outer diameter) as previously described.^[33]^ The scaffold microarchitecture (i.e., fiber morphology, diameter, and organization) was assessed from scanning electron microscopy (SEM) images (Quanta 600F; FEI, Hillsboro, OR) and the scaffold mechanical properties were characterized from biaxial stress–strain curves (CellScale Biomaterial Testing, Waterloo, Canada) as described below. Prior to cell seeding, the silicone-mounted scaffolds were sterilized by UV exposure (30 min per side), pre-wetted with water, and incubated overnight at 37 °C in complete medium (1:1 Roswell Park Memorial Institute-1640: advanced Dulbecco's modified Eagle (RPMI-1640:a-DMEM) medium (Gibco; ref A10491 and ref 124910) with 10 % fetal bovine serum (FBS, Greiner, Alphen aan den Rijn, The Netherlands), 1 % penicillin/streptomycin (P/S, Lonza, Basel, Switzerland; DE17-602E), 0.5 % GlutaMax (Gibco; ref 35050), and 0.25 mg mL^−1^ L-ascorbic acid 2-phosphate (Sigma; A8960)) to allow for protein adsorption.

### Human vena saphena cell culture

HVSCs were isolated from a single human donor after coronary by-pass surgery conforming to well-established protocols,^[66]^ in accordance to the Dutch advice for secondary-use material. The cells were cultured in complete a-DMEM (10 % FBS, 1 % GlutaMax, and 1 % P/S) in a standard culture incubator (37 °C, 5 % CO_2_) and passaged at 80 % confluency. Medium was refreshed every 3-4 days. For experiments, cells at passage 7 were used. Cell phenotype was assessed via qPCR.

### Peripheral blood mononuclear cell isolation and monocyte purification

Human peripheral blood mononuclear cells (hPBMCs) were isolated from two different buffy coats of healthy donors (purchased from Sanquin Blood Supply Foundation, Nijmegen, the Netherlands) using density gradient centrifugation on iso-osmotic medium (density: 1.077 g mL^−1^, Lymphoprep, Stemcell Technologies, Köln, Germany). In short, buffy coats were diluted in 0.6 % sodium citrate (Sigma-Aldrich, C7254) in phosphate-buffered saline (PBS), after which the solutions were carefully layered on top of an iso-osmotic medium (density 1.077 g mL^−1^, Lymphoprep, Axis Shield). The layered samples were centrifuged (21,000 rpm for 30 min) to extract the plasma and the erythrocytes from the white blood cell fraction. The hPBMCs were washed, re-suspended in freezing medium (RPMI-1640 supplemented with 20 % FBS, 1 % P/S, and 10 % Dimethyl sulfoxide (Merck Millipore)), and cryopreserved in liquid nitrogen until further use. For each donor, hPBMCs were characterized via flow cytometry (Guava easyCyte 6HT, Merck Millipore; **Table S2**). Prior to cell seeding, the hPBMC were thawed, resuspended in complete RPMI medium (10 % FBS, 1 % P/S), and counted. The monocytic cell fraction was isolated from the hPBMC fraction via negative selection using the MACS pan-monocyte isolation kit (Miltenyi Biotec) according to the supplier’s instructions. Monocyte purity (i.e., lymphocyte contamination expressed as the CD3/CD14 ratio), and the classical (CD14++ CD16-), non-classical (CD14+, CD16++), and intermediate (CD14++, CD16+) monocytes were quantified via flow cytometry (Table S2).^[67][68]^

### Cell seeding and in vitro culture

Human monocytes and HVSCs were seeded in a 2:1 ratio using bovine fibrin as a cell carrier.^[69]^ Prior to seeding, medium was removed from the scaffold and a suspension of bovine fibrinogen (10 mg mL^−1^, Sigma; F8630), bovine thrombin (10 IU mL^−1^, Sigma; T4648), and cells (30×10^6^ monocytes cm^−3^ and 15×10^6^ HVSCs cm^−3^) was carefully dripped and absorbed into the pre-wetted scaffold. Fibrin was allowed to polymerize for 60 min in an incubator (37 °C, 5 % CO_2_). Subsequently, the medium reservoirs were filled with 50 mL complete medium, and the cell-seeded constructs were mounted into the flow chamber and connected to the stretch pump and flow loop to start hemodynamic loading. Cyclic stretch was maintained at ~1.06 by adjusting the imposed pressure of the actuator. The applied stretches were monitored and quantified from the minimum and maximum outer diameter of the scaffolds, using time-lapse photography and custom Matlab scripts (Matlab R2017; The Mathworks, Natick, MA). Shear stress was maintained at ~1 Pa by tuning the flow rate magnitude to 60 mL/min (assuming a constant medium viscosity of 0.7·10^−3^ Pa s at 37 °C and *Q* = 61.14 · *τ*_*w*_ as previously described^[33]^). Applied wall shear stress was quantified from the flow magnitude during the course of the experiment and tuned by adjusting the pump pressure. Three times per week, 25 mL of medium was refreshed and supplemented to 50 mL. After 3 or 20 days of culture, tubular constructs were collected, sectioned according to a cutting scheme (**Figure S6**), and stored at 4 °C (after 15 min fixing in 3.7 % formaldehyde and 3×5 min washing in PBS) or −30 °C (after snap-freezing in liquid nitrogen) until further analysis. Additionally, supernatants at day 3 and day 20 were collected from the culture medium after centrifugation (300 g, 5 min). Supernatants were stored at −30 °C until further analysis.

### Scanning Electron Microscopy (SEM) analysis

Samples at day 3 and 20 were fixed in 2.5 % glutaraldehyde grade I (Sigma; G5882), washed in PBS (3×5 min) and dehydrated in a graded ethanol (EtOH) series, ranging from 50 % until 100 %. Before imaging, samples were dried and gold sputtered. Samples were visualized in high vacuum, using a 5 kV electron beam (Quanta 600F; FEI, Hillsboro, OR). Bare scaffold samples were directly visualized after drying in vacuum. Images were taken at different representative locations at multiple magnifications (50×, 500×, 1500×). The average fiber diameter of the bare scaffolds was quantified for 20 randomly chosen fibers in three representative images using the Image J software (U.S. National Institutes of Health, Bethesda, MD, USA).

### Immunohistochemistry analysis

Formalin-fixed samples were washed in PBS and permeabilized in 0.5 % Triton-X 100 in PBS for 30 min. Non-specific binding was blocked for 30 min using 5 % (w/v) goat serum in a PBS solution containing 1 % (w/v) BSA. Primary antibodies (in 10× diluted blocking solution) were incubated overnight at 4 °C (**Table S3**). After washing in PBS (3×5 min), the fluorescently labeled secondary antibodies (1:500 in 10× diluted block solution) were added for 1 h (Table S3), together with 4’,6-diamidino-2-phenylindole (DAPI, 1:500, Sigma) for localization of cell nuclei. Whole-mounts were kept in PBS and visualized with a confocal laser scanning microscope (Leica TCS SP5X with a 40×/1.1 HCX PL Apo CS lens). Cross-sections were mounted in mowiol and visualized with an inverted fluorescent microscope (Leica DMi8,with a 20×/0.4 or 40×/0.95 HC PL Fluotar lens).

### Fluorescence stainings

The cellular organization (i.e., actin cytoskeleton) at day 20 was visualized in formalin-fixed whole-mounts by phalloidin-Atto 488 (1:200, Sigma). The engineered tissue organization (i.e., collagen) was detected by CNA35-OG488 (1 μM^[70]^). After washing in PBS, cell nuclei in the actin-stained samples were labeled with propidium iodide (7 μM, Molecular probes; P3566). The stained samples were visualized with a confocal laser scanning microscope (Leica TCS SP5X with a 40×/1.1 HCX PL Apo CS lens). Actin and collagen distributions were quantified with in-house developed software as described elsewhere.^[33]^

### Histological analysis

Fibrillar collagen and glycosaminoglycans (GAGs) at day 20 were visualized in cross-sections with the Picrosirius red stain and Alcian blue stain, respectively. Images were acquired with a brightfield microscope (Zeiss Axio Observer Z1 with a 20×/0.8 Plan-Apochromat lens). The Picrosirius-stained samples were also imaged with polarized light illumination to study the birefringence of the collagen fibrils.

### Gene expression analysis

Prior to RNA extraction, samples were reduced to a fine powder with a micro-dismembrator and RLT buffer containing β-mercaptoethanol (Sigma; M3148) was added to further lyse the samples. Successively, RNA was extracted with the Qiagen RNeasy kit according to the manufacturer’s description. A 30 min DNAse incubation step (Qiagen; 74106) was performed to exclude genomic DNA contamination. RNA quantity and purity, and RNA integrity were assessed using, respectively, a spectrophotometer (NanoDrop, ND-1000, Isogen Life Science, The Netherlands) and gel electrophoresis. cDNA was synthesized in a thermal cycler (C1000 Touch, Bio-Rad) using a reaction solution consisting of RNA (100 or 200 ng) supplemented with 1 μL random primers (50 ng μL^−1^, Promega, C1181), 1 μL dNTPs (10 mM, Invitrogen), 2 μL 0.1M DTT, 4 μL 5× first strand buffer, 1 μL M-MLV Reverse Transcriptase (200 U μL^−1^, Invitrogen, 28025-013, Breda, the Netherlands), and RNAse-free ultra-pure water (ddH_2_O) up to a final volume of 11 μL. The reaction solution was exposed to the following thermal protocol: 65 °C (5 min), on ice (2 min) while adding the enzyme mixture, 37 °C (2 min), 25 °C (1 min), 37 °C (50 min), and 70 °C (15 min). Genomic contamination in the cDNA was assessed with glyceraldehydes-3-phosphate dehydrogenase (GAPDH) primers, conventional PCR, and gel electrophoresis.

After this check, qPCR was performed employing the gene-specific primer sequences listed in **Table S4**. GAPDH was identified as most stable and therefore selected as reference gene. Gene expression levels involved in inflammatory processes, cell phenotype, tissue formation, and remodeling were evaluated by adding 500 nM primer mix, 5 μl SYBR Green Supermix (Bio-Rad; 170-8886), and an additional 1.75 μL ddH_2_O to 3 μL cDNA (30× or 100× diluted). Solutions were exposed to the following thermal protocol: 95 °C (3 min), 40 cycles of 95 °C (20 s), 60 °C (20 s), and 72 °C (30 s), 95 °C (1 min), and 65 °C (1 min), completed with a melting curve measurement. C_t_ values were normalized for the reference gene and the 2^−ΔCt^ formula was applied to calculate relative expression levels.^[71]^

### ELISA

Secreted cytokines were quantified to obtain insight in the generated early (immune) profiles of the different experimental conditions (Table S1). Cytokines were quantified at the Multiplex core facility of the Laboratory for Translational Immunology of the University Medical Centre Utrecht (UMCU), the Netherlands, using a multiplex immunoassay built upon Luminex technology. In short, supernatants were incubated (1 h) with antibody-bound MagPlex microspheres (BioRad, Hercules, CA), succeeded by an incubation step with biotinylated antibodies (1 h), and a subsequent incubation with phycoerythrin-conjugated streptavidin (10 min) which was diluted in high performance ELISA (HPE) buffer (Sanquin). Data acquisition was executed using a FLEXMAP 3D system controlled with xPONENT 4.1 software (Luminex, Austin, TX), and analyzed by fitting a 5-parametric curve with Bio-Plex Manager software (version 6.1.1., Biorad).

### Biochemical assays

The engineered tissue constructs collected at day 20 were lyophilized, weighed, and reduced to a fine powder prior to determination of glycosaminoglycan (GAG), hydroxyproline (HYP), and DNA content. To obtain the powder, the constructs, together with 3 mm Ø beads (Sartorius, Goettingen, Germany), were added to Nalgene cryogenic vials (Thermo Scientific; 5000-0012), frozen in liquid nitrogen, and disrupted with a micro-dismembrator (Sartorius, 2×30 s at 3,000 rpm). Subsequently, 500 μL of digestion buffer (100 mM phosphate buffer (pH=6.5), 5 mM L-cysteine (C-1276), 5 mM ethylene-di-amine-tetra-acetic acid (EDTA, ED2SS), and 140 μg mL^−1^ papain (P4762), all from Sigma-Aldrich) was added, and the suspensions were transferred to new Eppendorf tubes for overnight digestion (16 h at 60 °C). Following digestion, samples were vortexed and centrifuged at 12,000 rpm for 10 min to precipitate scaffold remnants. The supernatant was used for DNA, GAG, and HYP quantification.

DNA content was quantified employing the Qubit dsDNA BR assay kit (Life Technologies, Carlsbad, California, USA) and the Qubit fluorometer (Life Technologies) according to the manufacturer’s description and normalized to total dry mass. GAG and HYP content were determined as indicators of ECM formation. GAG quantities were determined using a modified dimethyl methylene blue (DMMB) assay.^[72]^ Shark chondroitin sulfate (Sigma; C4348) was included as a standard. In short, 40 μL of the supernatants and prepared standards were transferred (in duplicate) to a 96-well plate. Subsequently, DMMB solution (150 μL) was added and absorbance (540 nm) was measured using a microplate reader (Synergy HTX; Biotek). To quantify HYP content, as a measure of mature collagen, digested samples were first hydrolyzed using 16 M sodium hydroxide (Merck; B1438798). Then, HYP content was quantified with a Chloramin-T assay, including trans-4-hydroxyproline as a reference (Sigma; H5534).^[73]^ GAG and HYP values were normalized to total dry mass to assess overall tissue formation.

### Mechanical analysis

The mechanical properties of bare scaffolds and constructs at day 20 were quantified using a biaxial tensile setup (CellScale Biomaterial Testing, Waterloo, Canada). Tests were performed in PBS at 37 °C. After 10 uniaxial stretch cycles (1.10) in the two orthogonal directions (i.e., axial and circumferential), the samples were equibiaxially stretched at a rate of 100 % min^−1^ while recording forces with a 1500 mN load cell. Graphite particles sprayed onto the sample’s surface facilitated optical strain analysis. Sample thickness was measured with a digital microscope (Keyence VHX-500FE). Stress–stretch curves were derived from the force and displacement measurements, assuming incompressibility and plane-stress conditions. Elastic moduli were computed as the slope of the stress–stretch curves at 10 % strain as a measure of stiffness.

### Statistical analysis

All data are reported as mean ± standard deviation unless stated otherwise. In addition, ELISA and qPCR data are displayed as heat maps using Heatmapper, a freely available web tool.^[74]^ Here, each cytokine/gene expression is transformed into a z-score (i.e., the number of standard deviations from the mean expression) and converted to a grayscale. To test for significant differences, the Kruskal-Wallis test with a Dunn’s multiple comparison test was performed using Prism (GraphPad, La Jolla, CA, USA). Prior to the statistical analysis, C_t_ values from the qPCR analysis were logarithmically transformed. Differences were considered significant for p-values < 0.05.

## Supporting Information

Supporting Information is available from the bioRχiv website or from the author.

## Acknowledgements

E.H. and T.W. contributed equally to this work. The authors would like to thank Prof. Anthony Weiss for critically assessing the manuscript, and Marina van Doeselaar and Marloes Janssen-van den Broek for their assistance in the sample analysis. This study (436001003) is financially supported by ZonMw as part of the LSH 2Treat program and the Dutch Kidney Foundation. We also gratefully acknowledge funding for the Gravitation Program “Materials-Driven Regeneration” by the Netherlands Organization for Scientific Research.

