## Supporting Information for "Human *in vitro* model of material-driven vascular regeneration reveals how cyclic stretch and shear stress differentially modulate inflammation and tissue formation"

**Table S1**. Genes and proteins analyzed via qPCR and Multiplex ELISA

| Protein | Symbol | Function | qPCR | ELISA | |
| --- | --- | --- | --- | --- | --- |
| Phenotypic markers |  |  |  |  |  |
| Monocyte chemoattractant protein 1 | MCP-1 | Chemotactic for monocytes/macrophages | x |  | x |
| Tumor necrosis factor alpha | TNF-α | Pro-inflammatory factor, stimulus for collagen production, inhibitor of elastogenesis | x |  | x |
| Interleukin 6 | IL-6 | Pro-inflammatory factor |  |  | x |
| Interleukin 10 | IL-10 | Anti-inflammatory cytokine, inhibitor of collagen production | x |  | x |
| Interleukin 13 | IL-13 | Anti-inflammatory factor, stimulus for collagen production |  |  | x |
| Alpha Smooth Muscle Actin | αSMA | Filament of the cytoskeleton involved in regulating cell shape, movement and involved in cell contractility | x |  |  |
| Smoothelin | SMTN | Constitutes part of the cytoskeleton and is found exclusively in contractile smooth muscle cells | x |  |  |
| Calponin | CNN1 | Protein that is involved in the modulation and regulation of smooth muscle cell contraction | x |  |  |
| S100 calcium binding protein A4 | S100A4 | Protein involved in the regulation of multiple cellular processes (e.g. cell cycle progression, differentiation, tubulin polymerization). Activated fibroblast, related to tissue remodeling. | x |  |  |
| Vimentin | VIM | Intermediate filament protein, part of the cytoskeleton, used as fibroblast marker | x |  |  |
| Tissue formation |  |  |  |  |  |
| Transforming growth  factor beta 1 | TGF-β1 | Anti-inflammatory factor; stimulus for collagen formation |  |  | x |
| Platelet derived growth factor- subunit BB | PDGF-BB | Stimulus for collagen formation and cell proliferation |  |  | x |
| Connective tissue growth factor | CTGF | Stimulus for collagen formation |  |  | x |
| Collagen Type I | Col Type I | Load bearing protein of the extracellular matrix | x |  |  |
| Collagen Type III | Col Type III | A fibrillar collagen; found frequently in association with type I collagen. | x |  |  |
| Lysyl Oxidase | LOX | Enzyme involved in collagen and elastin crosslinking | x |  |  |
| Elastin | ELN | Tropoelastin, one of the main components of the elastic fiber | x |  |  |
| Fibrillin-1 | FBN-1 | Extracellular matrix protein that provides structural support for elastic fibril formation | x |  |  |
| Fibrillin-2 | FBN-2 | Extracellular matrix protein that provides structural support for elastic fibril formation | x |  |  |
| Decorin | DCN | Proteoglycan important in collagen fibril assembly | x |  |  |
| Versican | VCAN | Proteoglycan important for cell adhesion, proliferation, differentiation and migration. Proven important for elastic network formation | x |  |  |
| Fibronectin | FN | Extracellular adhesion protein involved in adhesion, growth, migration, and differentiation | x |  |  |
| Remodeling |  |  |  |  |  |
| Matrix metalloproteinase 9 | MMP-9 | Anti-inflammatory factor involved in extra-cellular breakdown and remodeling |  |  | x |
| Matrix metalloproteinase 1 | MMP-1 | Extra-cellular breakdown and remodeling | x |  | x |
| Matrix metalloproteinase 2 | MMP-2 | Extra-cellular breakdown and remodeling | x |  |  |
| Metallopeptidase inhibitor 1 | TIMP-1 | Inhibitor of MMPs | x |  | x |
| Metallopeptidase inhibitor 2 | TIMP-2 | Inhibitor of MMPs | x |  |  |

**Table S2**. Immune cell characterization

|  |  | Mean ± SD d706 | Mean ± SD d669 | Physiological range (min-max) |
| --- | --- | --- | --- | --- |
| PBMCs | | | | |
| Monocytes  (% of CD3/CD14+) |  | 29.2 ± 1.2 | 32.4 ± 4.0 | 7.9 - 37.5 |
| (% of MON1/2/3) | MON1 | 83.7 ± 2.6 | 79.1 ± 2.1 | 67.2 - 94.2 |
|  | MON2 | 7.4 ± 2.0 | 8.5 ± 0.7 | 1.2 - 10.2 |
|  | MON3 | 8.9 ± 0.6 | 12.4 ± 2.8 | 4.6 - 22.6 |
| Lymphocytes  (% of CD3/CD14+) |  | 70.8 ± 1.2 | 67.6 ± 4.0 | 62.5 - 92.1 |
| (% of lymphocyte cloud) | Th-cells | 37.4 ± 0.4 | 39.1 ± 0.6 | 18.0 - 58.5 |
|  | Tc-cells | 20.5 ± 0.4 | 17.4 ± 0.2 | 17.1 - 39.1 |
|  | DP T-cells | 0.8 ± 0.1 | 0.8 ± 0.2 | 0.6 - 2.5 |
|  | CD4/CD8 ratio | 1.8 ± 0.1 | 2.3 ± 0.1 | 0.5 - 2.8 |
| After monocyte isolation | | | | |
| Monocytes  (% of CD3/CD14+) |  | 91.0 ± 14.4 | 88.5 ± 15.3 |  |
| (% of MON1/2/3) | MON1 | 77.0 ± 5.0 | 75.8 ± 4.2 |  |
|  | MON2 | 6.2 ± 0.7 | 10.5 ± 2.2 |  |
|  | MON3 | 16.9 ± 4.5 | 13.7 ± 2.2 |  |
| Lymphocytes  (% of CD3/CD14+) |  | 9.0 ± 14.4 | 11.5 ± 15.3 |  |
| Before Isolation | CD3/CD14 ratio | 2.4 ± 0.1 | 2.1 ± 0.4 |  |
| After isolation | CD3/CD14 ratio | 0.1 ± 0.2 | 0.2 ± 0.2 |  |

* Abbreviations: monocytes (MON), helper T-cells (Th-cells), cytolytic T-cells (Tc-cells) and double positive T-cells (DP T-cells)

The hPBMCs of both donors were characterized via flow cytometry (FACs, Guava easyCyte 6HT, Merck Millipore) using the conjugated monoclonal antibodies anti-CD14 (FITC, AbD Serotec), anti-CD16 (Alexa 647, AbD Serotec), CD3 (PerCP/Cy5.5, BioLegend), CD4 (FITC, Abcam), and CD8 (PE, Abcam). In brief, non-specific binding was blocked for 10 min on ice with 2 % bovine serum albumin (BSA, Roche) or 2 % human serum (Lonza, for the CD16 Ab, to prevent unspecific binding) in PBS. Subsequently, cells were centrifuged (1,400 rpm, 4 °C, 5 min) and the blocking solution was removed. The conjugated monoclonal antibodies were diluted in 0.5 % BSA in PBS to the optimal working concentrations and incubated for 1 h at 4 °C in the dark, followed by a PBS washing step on ice (3×5 min). Cells were either directly analyzed or fixed in 3.7 % formaldehyde for 10 min (+3×5 min PBS washing step) and analyzed the next day. Prior to FACs analysis, cells were well suspended in 0.5 % BSA in PBS to remove cell clumps. Gating and additional data analyses were performed using Guava Express Pro Software. Cell populations were quantified starting from the initial monocyte and lymphocyte cloud (defined as hPBMC population) that were identified in the forward scatter - side scatter plots. Within this hPBMC gate, monocytes were identified as Monocyte 1 (MON 1; CD14+/CD16-), Monocyte 2 (MON 2; CD14+/CD16+), and Monocyte 3 (MON 3; CD14 dim/CD16+).^[67]^ Lymphocytes were further characterized to differentiate between helper T-cells (Th-cells; CD3+/CD4+/CD8-), cytolytic T-cells (Tc-cells; CD3+/CD4-/CD8+), and double positive T-cells (DP T-cells; CD3+/CD4+/CD8+).^[68]^ Overall, all cell quantities of the specified cell types fell within the physiological values of healthy adults. Further cell purification resulted in an 80-90 % pure monocyte population (Table S2).

**Table S3**. Markers selected for immunohistochemistry for the localization of monocytes (CD45-positive) and HVSCs (vimentin-positive), proliferation (KI67), and matrix content (collagen type I/III)

| Target antigen | Vendor or Source | Working concentration | Comments |
| --- | --- | --- | --- |
| Primary antibodies | | | |
| CD45, mouse IgG1 | Abcam | 1:1000 | Cell  (whole-mount (day 3) / cross-section (day 20)) |
| Vimentin, mouse IgM | Abcam | 1:2000 |  |
| KI67,rabbit IgG | ThermoScientific | 1:200 |  |
| collagen I, mouse IgG1 | Sigma | 1:200 | Matrix  (cross-section) |
| collagen III, rabbit IgG | Abcam | 1:250 |  |
| Secondary antibodies | | | |
| goat anti-mouse IgG1-555 | Molecular Probes | 1:500 | Cell  (whole-mount (day 3) / cross-section (day 20)) |
| goat anti-mouse IgM-647 | Jackson ImmunoResearch | 1:500 |  |
| goat anti-rabbit IgG-488 | Molecular Probes | 1:500 |  |
| goat anti-mouse IgG-488 | Molecular Probes | 1:500 | Matrix  (cross-section) |
| goat anti-rabbit IgG-647 | Molecular Probes | 1:500 |  |

**Table S4**. Primers for gene expression analysis

| Primer | Symbol | Accession number | Primer Sequence (‘5-‘3) |
| --- | --- | --- | --- |
| Phenotypic markers | | | |
| α smooth muscle actin | ACTA2 | NM_001613.1 | FW: CGTGTTGCCCCTGAAGAGCAT  RV: ACCGCCTGGATAGCCACATACA |
| Smoothelin | SMTN | NM_134270 | FW: CAGCCCAGAACCGAGAGTC  RV: AGCAGCCATAGGAGAATCAGAT |
| Calponin | CNN1 | NM_001299.5 | FW:TGAAGTACGCAGAGAAGCAG RV:CAGCTTGGGGTCGTAGAG |
| Vimentin | VIM | NM_003380 | FW: AAGACCTGCTCAATGTTAAGATC  RV: CTGCTCTCCTCGCCTTCC |
| S100 Calcium Binding Protein A4 | S100A4 | NM_002961 | FW:TCTTTCTTGGTTTGATCCTGACT RV: AGTTCTGACTTGTTGAGCTTGA |
| Tissue formation | | | |
| Transforming growth  factor, beta 1 | TGFB1 | NM_000660 | FW: GCAACAATTCCTGGCGATACCTC  RV: AGTTCTTCTCCGTGGAGCTGAAG |
| Collagen type I | COL1A1 | NM_000088 | FW: AATCACCTGCGTACAGAACGG  RV:TCGTCACAGATCACGTCATCG |
| Collagen type III | COL3A3 | NM_000090 | FW: ATCTTGGTCAGTCCTATGC  RV: TGGAATTTCTGGGTTGGG |
| Lysyl Oxidase | LOX | NM_002317.3 | FW: CCTGGCTGTTATGATAC RV: GAGGCATACGCATGATG |
| Fibronectin | FBN | NM_001306129.1 | FW: AAGACCAGCAGAGGCATAAGG RV: CACTCATCTCCAACGGCATAATG |
| Elastin | ELN | NM_000501.3 | FW: CTGGAATTGGAGGCATCG  RV: TCCTGGGACACCAACTAC |
| Fibrillin 1 | FBN1 | NM_00138 | FW:TGTTGGTTTGTGAAGATATTG RV: GTGGAGGTGAAGCGGTAG |
| Fibrillin 2 | FBN2 | NM_001999 | FW:ATCCCTGTGAGATGTGTC RV: TTCCTCCTTGGCATATCC |
| Decorin | DCN | NM_133503 | FW:TGCAGCTAGCCTGAAAGGAC RV: TTGGCCAGAGAGCCATTGTC |
| Versican | VCAN | NM_004385 | FW:GGCACCTGTTATCCTACTGAAA RV: ACACAAGTGGCTCCATTACG |
| Fibronectin | FN1 | NM_001306129.1 | FW: AAGACCAGCAGAGGCATAAGG  RF: CACTCATCTCCAACGGCATAATG |
| Remodeling | | | |
| Matrix Metalloproteinase 1 | MMP1 | NM_001145938.1 | FW:CGCACAAATCCCTTCTACCC RV: CTGTCGGCAAATTCGTAAGC |
| Matrix Metalloproteinase 2 | MMP2 | NM_001127891 | FW: ATGACAGCTGCACCACTGAG  RV: ATTTGTTGCCCAGGAAAGTG |
| Metallopeptidase inhibitor 1 | TIMP1 | NM_003254.2 | FW: TGACATCCGGTTCGTCTACA  RV: TGCAGTTTTCCAGCAATGAG |
| Metallopeptidase inhibitor 2 | TIMP2 | NM_003255.4 | FW:GGAGGAATCGGTGAGGTC RV: AACAGGCAAGAACAATGG |
| Inflammatory marker | | | |
| Monocyte chemoattractant  protein 1 | MCP1 | NM_002982 | FW: CAGCCAGATGCAATCAATGCC  RV: TGGAATCCTGAACCCACTTCT |

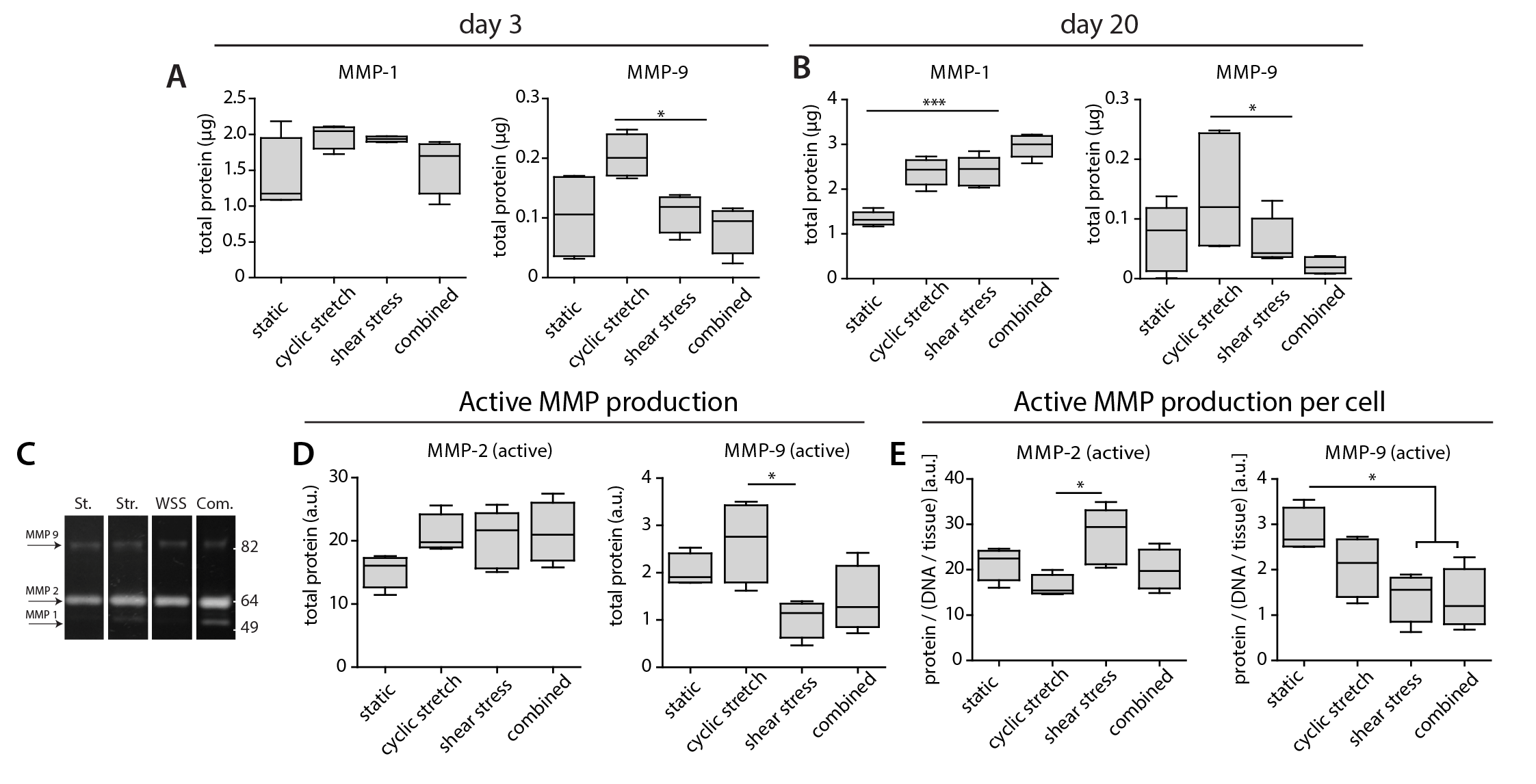

**Figure S1**. Matrix remodeling. A, B) Total matrix metalloproteinase (MMP) production at day 3 (*n* = 4 per group) and day 20 (*n* ≥ 5 per group) measured by ELISA. C) Gelatin zymogram gel for the detection of active MMPs at day 20. D, E) Quantification of band intensities for total active MMP production and cellular active MMP production (i.e., normalized to DNA content at day 20, *n* ≥ 5 per group) measured by gelatin zymography (* *p* < 0.05, *** *p* < 0.001). Abbreviations: matrix metalloproteinase (MMP).

Supernatant collected at day 20 was analyzed by gelatin zymography for the detection of MMP-2 and MMP-9. First, proteins were separated with gel electrophoresis in a 10 % acrylamide gel containing 0.6 mg mL^-1^ gelatin (Sigma; G-8150). After protein separation, gels were washed with 2.5% Triton X-100 (2× 30 min at 37 °C) and incubated with digestion buffer (50 mM Tris containing 0.05 % (w/v) CaCl_2_, pH 8.5) during 16 h at 37 °C. After digestion, the gels were stained with 0.1 % Brilliant Blue R 250 and de-stained with 10 % acetic acid and 4 % methanol. Band intensities were quantified with ImageJ (U.S. National Institutes of Health, Bethesda, MD, USA). Intensities were corrected for total culture medium volumes and DNA content. The zymography results clearly show that more MMP-2 than MMP-9 is produced, and that both types of MMPs are active (Figure S2C-E). In addition, in line with the ELISA data (Figure S2A, B), MMP-9 production seems to be reduced in the presence of hemodynamic loads. For MMP-2, however, the effect is less obvious. For the detection of MMP-1, the collagenase that was clearly enhanced with hemodynamic loading (Figure S2A, B), it is recommended to perform collagen zymography.

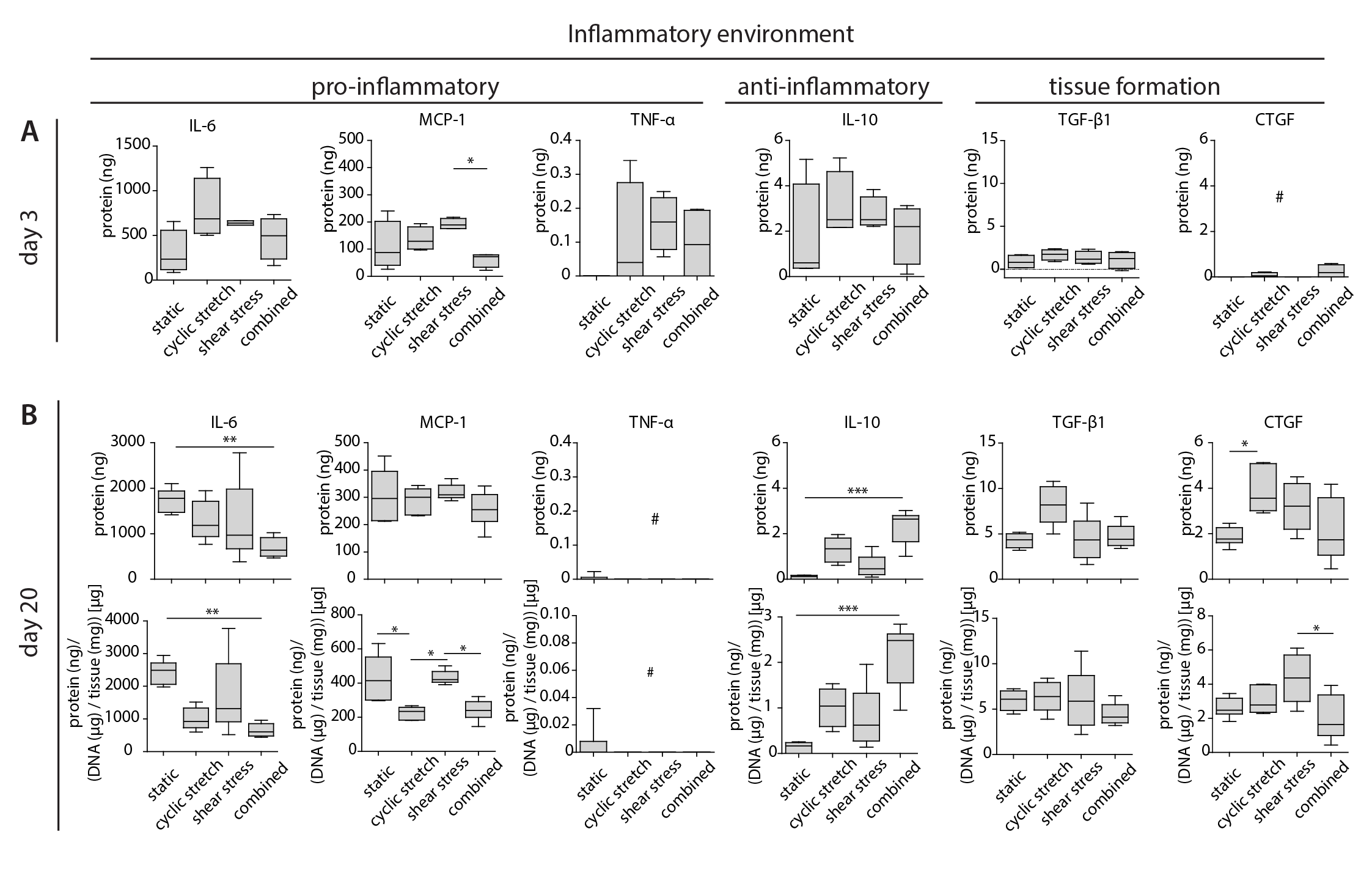

**Figure S2**. Cytokine secretion in the medium. A) Total cytokine secretion at day 3 (n = 4 per group). B) Total cytokine secretion at day 20 (*top, n* ≥ 5 per group), and normalized to DNA content at day 20 (*bottom*) (* *p* < 0.05, ** *p* < 0.01, *** *p* < 0.001; TNF-α was only detected in the mechanical loaded groups at day 3 (*n* ≥ 2); # close and below the detection limit). Abbreviations: interleukin (IL), monocyte chemoattractant protein 1 (MCP-1), tumor necrosis factor alpha (TNF-α), transforming growth factor beta (TGF-β1), connective tissue growth factor (CTGF).

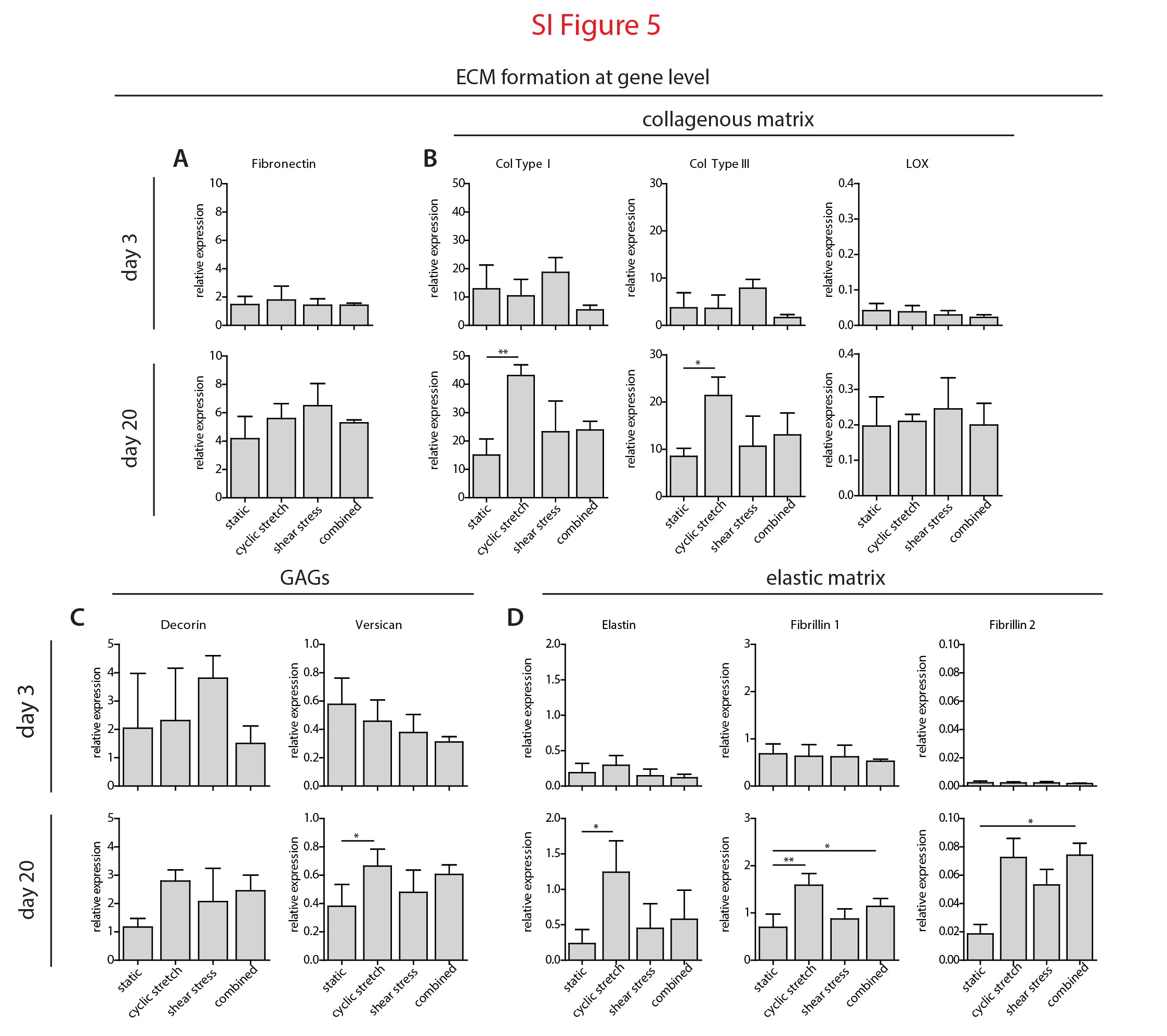

**Figure S3**. Matrix growth at day 3 and day 20. Gene expression (normalized to GAPDH) of A) fibronectin, B) collageneous matrix markers, C) glycosaminoglycan (GAG) markers, and D) elastic matrix markers. *n* ≥ 3 per group (day 3) and *n* ≥ 4 per group (day 20), except for LOX at day 3 (combined *n* = 2) and fibrilin-2 at day 20 (static *n* = 3 and cyclic stretch *n* = 2). * *p* < 0.05, ** *p* < 0.01. Abbreviations: col type (collagen type), lysyl oxidase (LOX).

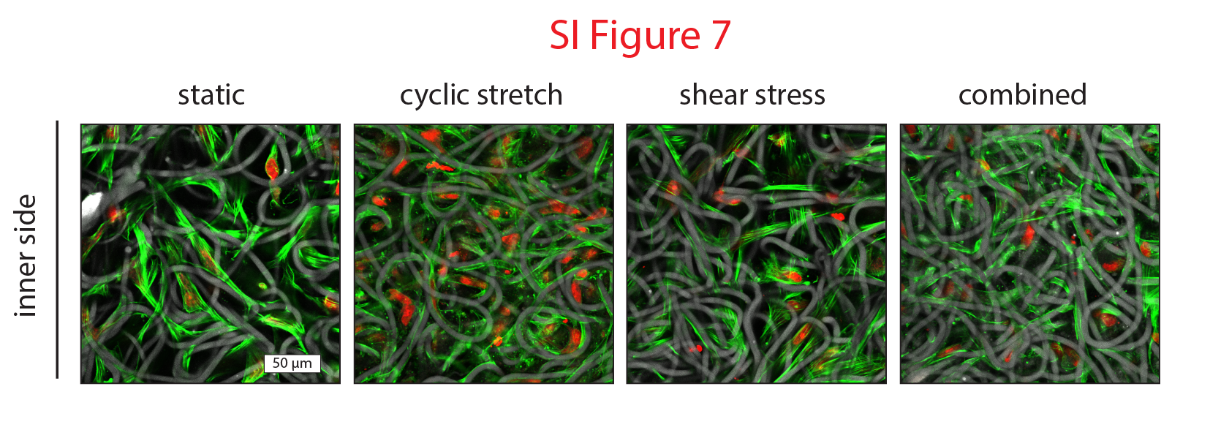

**Figure S4** Actin organization at day 20. Visualization of the actin fibers at the inner side of the constructs.

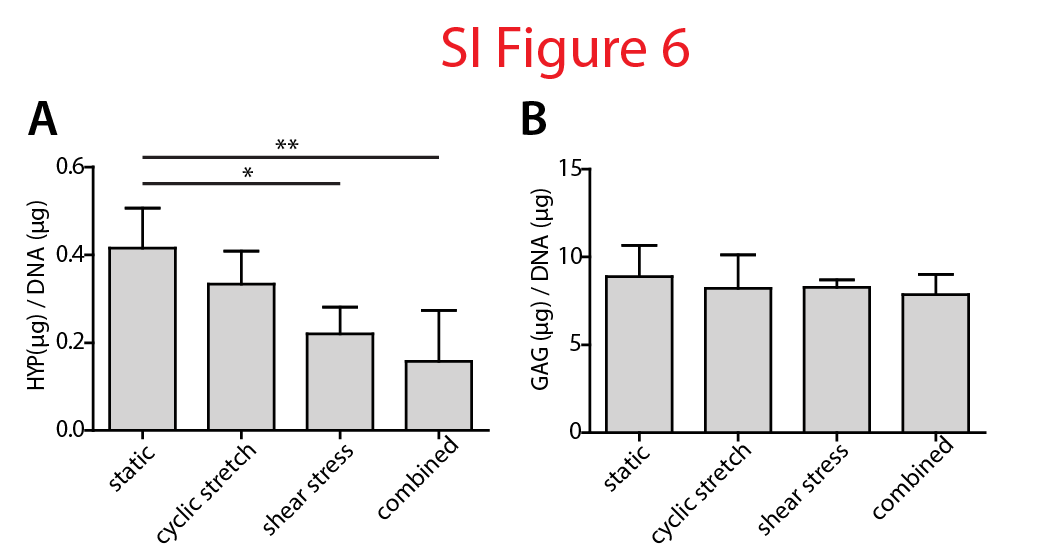

**Figure S5**. Biochemical assays. A) Hydroxyproline and B) glycosaminoglycan content normalized to DNA content, *n* ≥ 5 per group (* *p* < 0.05, ** *p* < 0.01). Abbreviations: hydroxyproline (HYP), glycosaminoglycans (GAGs).

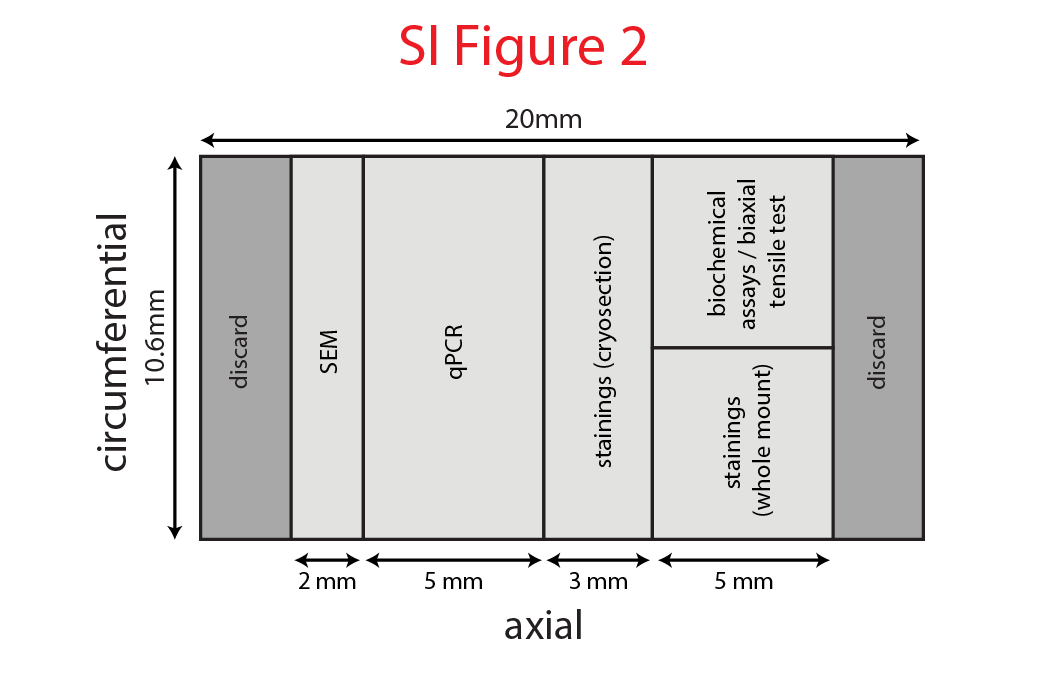

**Figure S6**. Experimental cutting scheme. Cutting scheme used for day 3 and at day 20 (N.B. at day 3, samples for biochemical assays / biaxial tensile test were pooled with samples for qPCR to isolate RNA).
